# Preliminary Study on the Mechanisms of Cytokinin and Its Receptor Binding Diversity

**DOI:** 10.1101/2025.08.05.668571

**Authors:** Xiang Bao, Chuan-Miao Zhou, Jia-Wei Wang

**Affiliations:** University College London, London WC1E 6BT, United Kingdom; CAS Center for Excellence in Molecular Plant Sciences (CEMPS), Institute of Plant Physiology and Ecology (SIPPE), Chinese Academy of Sciences (CAS), Shanghai 200032, China

**Keywords:** Cytokinins, Microscale Thermophoresis (MST), Molecular docking, Cytokinin receptor

## Abstract

Cytokinins are key regulators of plant growth and development, directly influencing crop yield. However, the quantitative basis of cytokinin receptor–ligand interactions remain poorly understood. This study aimed to quantitatively assess the binding affinities of cytokinin receptors various cytokinins, including 6-benzylaminopurine (6-BA), *N^6^*-isopentenyladenine (2iP), and trans-Zeatin (tZ) and explore cross-species conservation in cytokinin recognition. Microscale Thermophoresis (MST) assays revealed that 6-BA exhibits a higher binding affinity to the CHASE domain of the cytokinin receptor *Arabidopsis* HK4 (AHK4^CD^) compared to 2iP. Molecular docking and phylogenetic analyses further confirmed this finding and identified conserved key residues responsible for cytokinin binding across diverse plant species. Intriugingly, despite differences in pocket structure and binding energy, the cytokinin receptor in the bryophyte *Marchantia polymorpha* retains the capacity to bind cytokinins and respond to exogenous applications. Similarly, one cytokinin receptor in the monocots *Oryza sativa* exhibits distinct features in its cytokinin-binding pocket while still maintaining its ability to bind cytokinins. Our results thus provide fresh quantitative evidence for cytokinin receptor selectivity and highlight the structural plasticity of cytokinin perception, offering a foundation for designing optimized cytokinins or their receptors in crops.

## INTRODUCTION

Cytokinins are essential plant hormones that regulate key developmental processes, including cell proliferation, tissue patterning, and organogenesis (Kieffer et al., 2010). They are widely applied in agriculture and scientific research, such as tissue culture and crop improvement (Kieber & Schaller, 2018; Hnatuszko-Konka et al., 2021). Chemically, cytokinins are adenine derivatives substituted at the *N^6^* position, with representative examples including 6-benzylaminopurine (6-BA) and *N^6^*- isopentenyladenine (2iP) (Figure 1).

**Figure 1.**
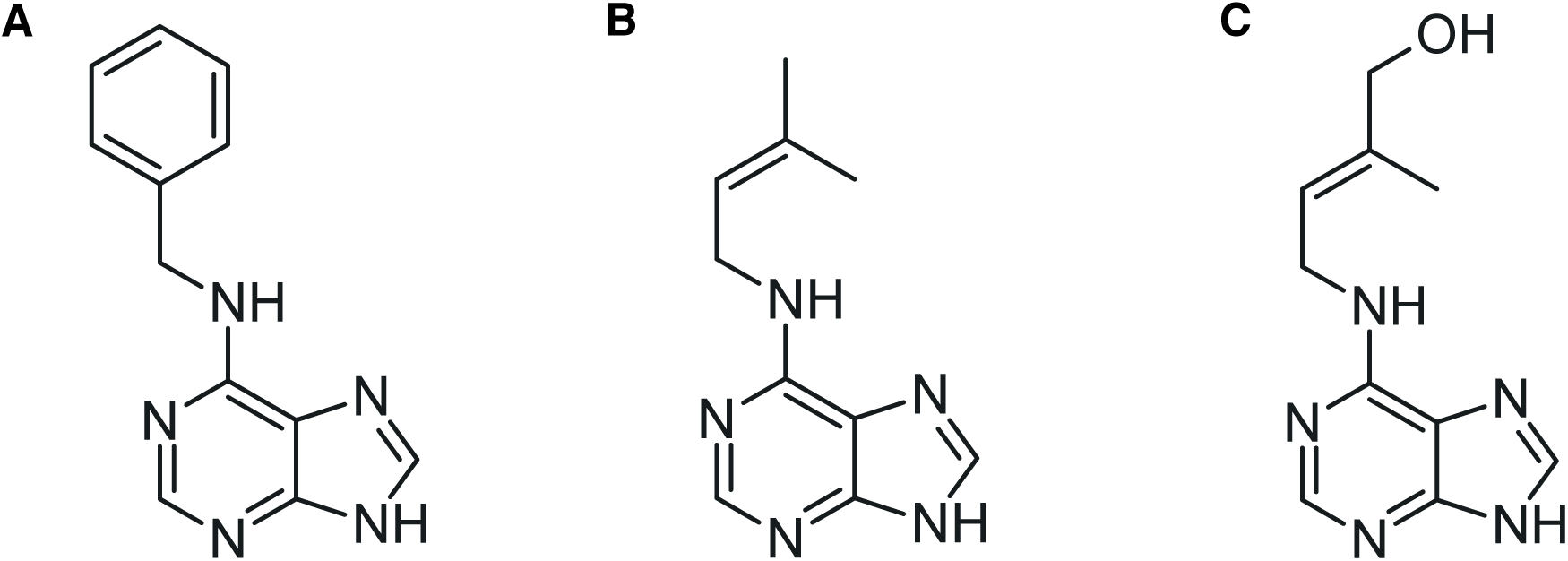
Chemical structures of three representative cytokinins. The three cytokinins differ in their *N⁶*-side chains: 6-BA (**A**) carries a benzyl group (–CH₂–C₆H₅), 2iP (**B**) has an isopentenyl group (–CH₂–CH=C(CH₃)₂), and tZ (**C**) has a hydroxylated isopentenyl group (–CH₂–CH(OH)–CH=C(CH₃)₂). The chemical structures were generated using ChemAxon MarvinSketch (https://chemaxon.com/) based on molecular information retrieved from PubChem (CID: 62389, 92180, and 449093).

Cytokinin perception in *Arabidopsis thaliana* is mediated by membrane-bound histidine kinase receptors (HKs), mainly AHK2, AHK3, and AHK4 (El-Showk et al., 2013). These receptors contain an extracellular CHASE (cyclases, histidine kinase associated sensory extracellular) domain (CD), which directly recognizes and binds cytokinins, thereby triggering a conserved two-component signaling (TCS) cascade involving a histidine kinase and a response regulator (Hothorn et al., 2011; Bauer et al., 2013; Kieber & Schaller, 2018). As the first and most critical step in cytokinin signaling, receptor–ligand interactions determine the specificity and strength of downstream responses (Romanov et al., 2006). Structural features within the CD largely define ligand selectivity, and even single amino acid substitutions, such as the Thr301Ile mutation in AHK4, can change cytokinin sensitivity (Yamada et al., 2001).

Understanding the structural and quantitative basis of cytokinin–receptor interactions is essential for explaining differences in cytokinin biological activity and for guiding the design of cytokinins with improved stability, efficiency, and selectivity. Such insights can also inform receptor engineering strategies to enhance ligand-binding affinity. Together, these approaches hold great potential for agricultural applications and plant science research (Kieber & Schaller, 2018; Hnatuszko-Konka et al., 2021).

Despite extensive studies on cytokinin signaling, the quantitative characterization of receptor–ligand binding remains limited. Most previous research has focused on downstream transcriptional responses, often neglecting the quantification of primary ligand–receptor interaction (Wulfetange et al., 2011; Bauer et al., 2013). Although AHK4 has been reported to bind multiple cytokinin variants, its precise binding affinities to different ligands have not been systematically evaluated (Hothorn et al., 2011). A detailed quantitative analysis of cytokinin binding is therefore crucial to link structural determinants with receptor selectivity and to guide the development of more effective cytokinin analogs.

In this study, we aimed to provide a quantitative assessment of cytokinin–receptor interactions to address the current lack of direct biophysical characterization. Microscale Thermophoresis (MST) (Jerabek-Willemsen et al., 2014) was employed as an *in vitro* system to determine the binding affinities of the AHK4 CHASE domain (AHK4^CD^) to two representative cytokinins, 6-BA and 2iP. To extend these findings beyond *Arabidopsis*, molecular docking was performed on homologous cytokinin receptors from *Marchantia polymorpha* and *Oryza sativa* with three major cytokinins (6-BA, 2iP, and trans-Zeatin (tZ)), aiming to explore cross-species conservation and divergence in cytokinin recognition.

## RESULTS

### Expression and Purification of 6×His-GFP-tagged AHK4^CD^

To investigate the ligand-binding affinity of cytokinin receptors *in vitro*, we cloned the AHK4^CD^ and fused it with a Green Fluorescent Protein (GFP) tag (AHK4^CD^-GFP) at the C-terminal. This design enables fluorescence-based MST detection and eliminates the need for external labeling reagents (Jerabek-Willemsen et al., 2014). The structure of the fusion protein was inferred using AlphaFold3 (Roy & Al-Hashimi, 2024) to confirm that AHK4^CD^ is structurally separated with GFP (Figure 2A). Additionally, an N-terminal 6×His tag is linked via a short, non-functional linker region and was used as a high affinity tag for protein purification (6×His-AHK4^CD^-GFP). The predicted molecular weight of the fusion protein is approximately 58.7 kDa. A 6×His-GFP fusion protein was used as a control in MST assays to exclude non-specific binding of cytokinins to the GFP tag (Figure 2B). The predicted molecular weight of the fusion protein is approximately 28.0 kDa.

**Figure 2.**
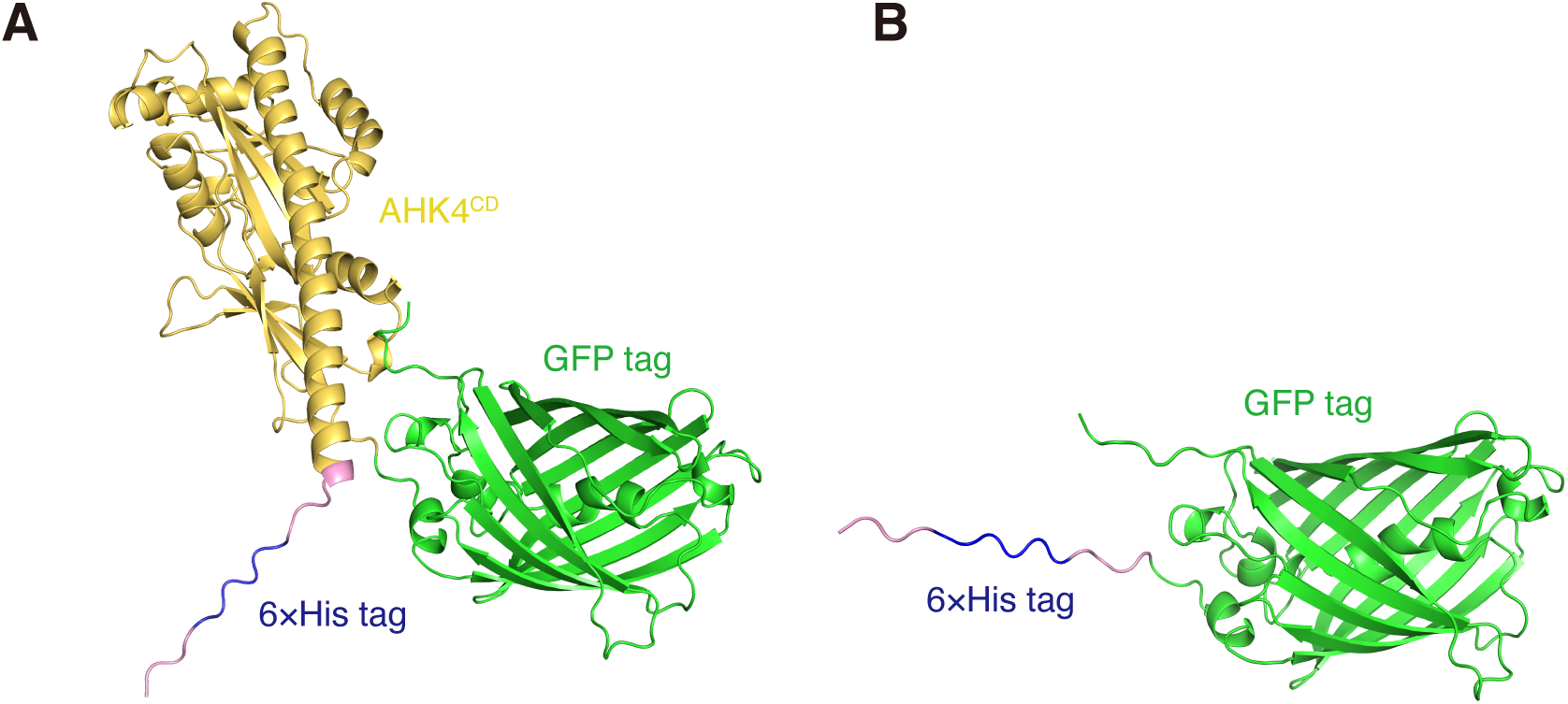
Predicted structures of the 6×His-AHK4^CD^-GFP and 6×His-GFP fusion proteins. **(A)** The structural model shows the AHK4^CD^ (yellow) fused with GFP (green) at the C-terminal and a 6×His tag (blue) at the N-terminal. The linker region between 6×His tag and AHK4^CD^ is highlighted in pink. **(B)** The structural model shows the GFP (green) protein fused with a 6×His tag (blue) at the N-terminal. This protein is used as a control for MST assays.

To confirm successful expression and purification of the recombinant proteins, Sodium Dodecyl Sulphate-Polyacrylamide Gel Electrophoresis (SDS-PAGE) analysis was performed following Ni^2+^- NTA affinity purification. A distinct band corresponding to ∼58 kDa was observed for the 6×His-AHK4^CD^-GFP fusion protein, consistent with the expected molecular weight. The control 6×His-GFP protein showed a distinct band at ∼28 kDa (Figure 3A). Minimal background bands indicated good purity for both preparations. In addition, the visible green fluorescence of the purified fusion protein further confirmed correct folding and stability of the GFP-tag (Figure 3B).

**Figure 3.**
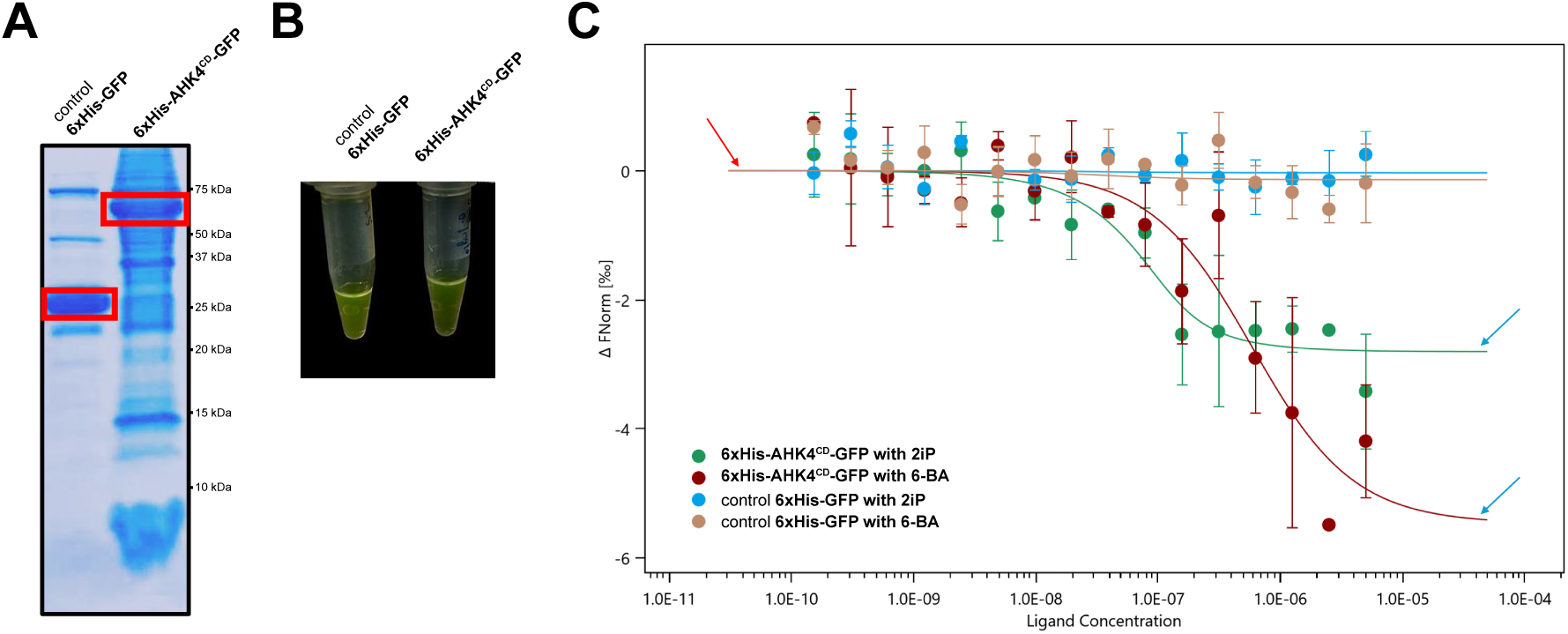
MST binding assays of 6×His-AHK4^CD^-GFP to 6-BA and 2iP. **(A)** Coomassie blue-stained SDS-PAGE showing the purification of 6×His-AHK4^CD^-GFP and 6×His-GFP fusion proteins. **(B)** Visualization of the fusion proteins under low-light conditions. **(C)** MST binding curves of 6×His-AHK4^CD^-GFP to 6-BA and 2iP. *Kd* values for 6×His-AHK4^CD^-GFP and the ligands 6-BA or 2iP binding were determined by MST. The X-axis represents the logarithm of ligand concentration (nM), while the Y-axis shows the normalized MST fluorescence response, reflecting the changes in thermophoretic mobility due to ligand binding. Error bars represent standard deviation (SD) from three independent biological replicates (*n* = 3).

### MST Analysis of Cytokinin Binding Affinity of AHK4^CD^

We then employed MST (Jerabek-Willemsen et al., 2014) to assess the binding affinities of the purified 6×His-AHK4^CD^-GFP proteins for two commonly studied cytokinins, 6-BA and 2iP (Figure 1). Specifically, we fixed the concentration of 6×His-AHK4^CD^-GFP proteins (target) at 100 nM, while the cytokinins (ligands) were serially diluted from 5 µM to 0.000153 µM.

The dissociation constant (*Kd*) for 6-BA was found to be 27.66 ± 20.73 nM, which is significantly lower than that for 2iP (540.21 ± 261.97 nM), showing a substantially higher binding affinity of the receptor for 6-BA (Figure 3C). Importantly, both datasets exhibited good signal-to-noise ratios (>8), underscoring the reliability of the curve fitting (Figure 3C).

In the experimental groups for 6-BA and 2iP, the MST fluorescence response changed with varying ligand concentrations, resulting in two distinct curves. In each curve, the left plateau (red arrow) corresponds to the unbound 6×His-AHK4^CD^-GFP, while the right plateau (blue arrows) indicates the bound target-ligand complex.

Control experiments performed with 6×His-GFP in the presence of either 6-BA (brown curve) or 2iP (blue curve) showed no significant changes in MST fluorescence response (i.e., both lines remained flat), confirming that the observed binding responses is specifically from the AHK4^CD^ rather than from the GFP tag.

### Structural Alignment of the CDs of AHKs

Given the well-established role of AHK4^CD^ in cytokinin perception, we utilized AlphaFold3 (Roy & Al- Hashimi, 2024) to predict and comparatively analyze the CD structures of the other two major cytokinin receptors, AHK2 and AHK3 in *A. thaliana*. Structural alignment revealed a high degree of similarity, with low root mean square deviations (RMSDs) of 0.696 Å for AHK2–AHK4 and 0.816 Å for AHK3– AHK4 (Figures 4A and 4B). These results support the structural conservation of the ligand-binding regions among the three AHK receptors, highlighting the use of AHK4 as a structurally representative model for the cytokinin receptor family in *Arabidopsis*.

**Figure 4.**
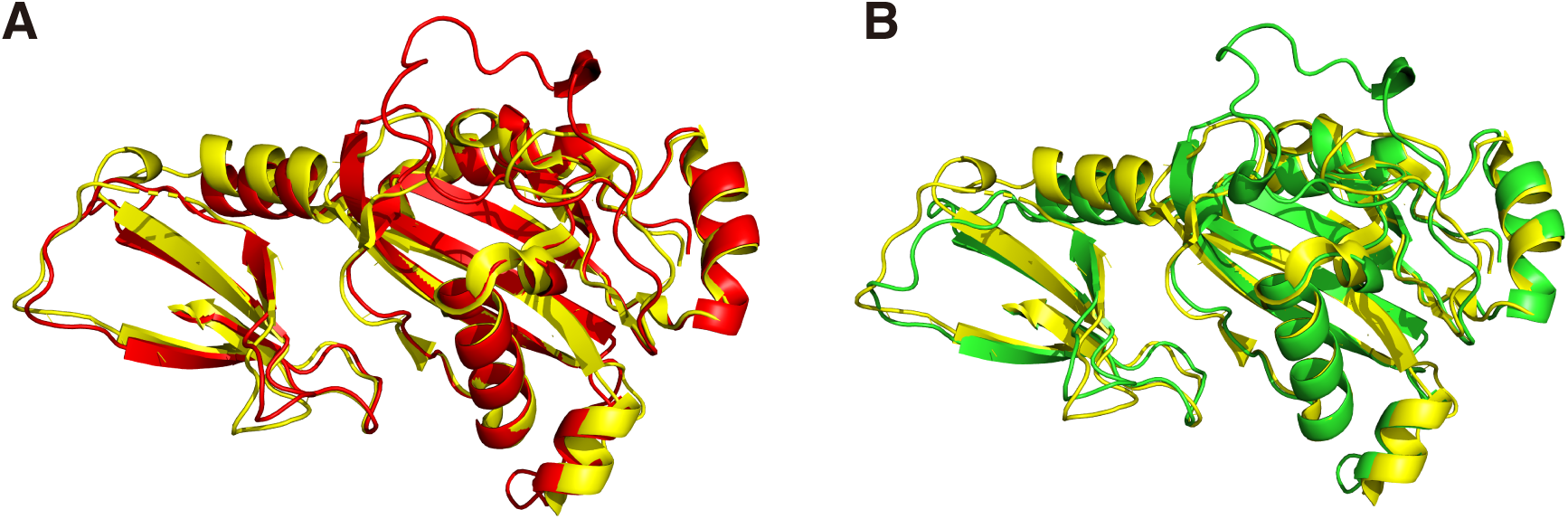
Structural alignment of AHK2^CD^, AHK3^CD^, and AHK4^CD^. **(A)** Ribbon diagram showing the structural alignment of the CD structures of AHK2 (**A**, red) and AHK3 (**B**, green) with that of AHK4 (yellow, PDB: 3T4J). The structures are aligned using PyMOL (The PyMOL Molecular Graphics System, Version 3.0 Schrödinger, LLC.).

### Probing Diverse Cytokinin Recognition Patterns through Molecular Docking Analyses

To explore whether cytokinin receptors from different plants exhibit distinct cytokinin recognition patterns, molecular docking was performed for the CD for four receptor proteins: Os02t0738400 (OHK6^CD^) and its homolog BGIOSGA009018 (B8AI44^CD^) from *O. sativa subsp. japonica* and *O. sativa subsp. indica*, respectively, as representatives for monocots, along with HK1 (MpHK1^CD^) and HK2 (MpHK2^CD^) from *M. polymorpha* as a representative for bryophytes. Each receptor was docked with 6- BA, 2iP and tZ. Receptor structures were predicted by AlphaFold3 (Roy & Al-Hashimi, 2024), ligand– receptor interactions were predicted using AutoDock (Morris et al., 2009; Goodsell et al., 2020), which identifies the lowest-energy binding pose by simulating ligand flexibility and scoring interaction energies.

Molecular docking of three cytokinins revealed consistent binding energy patterns across tested receptors. In all cases, 6-BA exhibited the most negative binding energy values, followed by 2iP, while tZ showed the least negative values (Figures 5 and 6; Supplementary Table 1). However, visualization of docking positions indicated that binding energy did not correlate with the number of direct hydrogen bonds (H-bonds) formed between ligand and receptor. For instance, several 6-BA docking models displayed strong binding despite forming only one H-bond.

**Figure 5.**
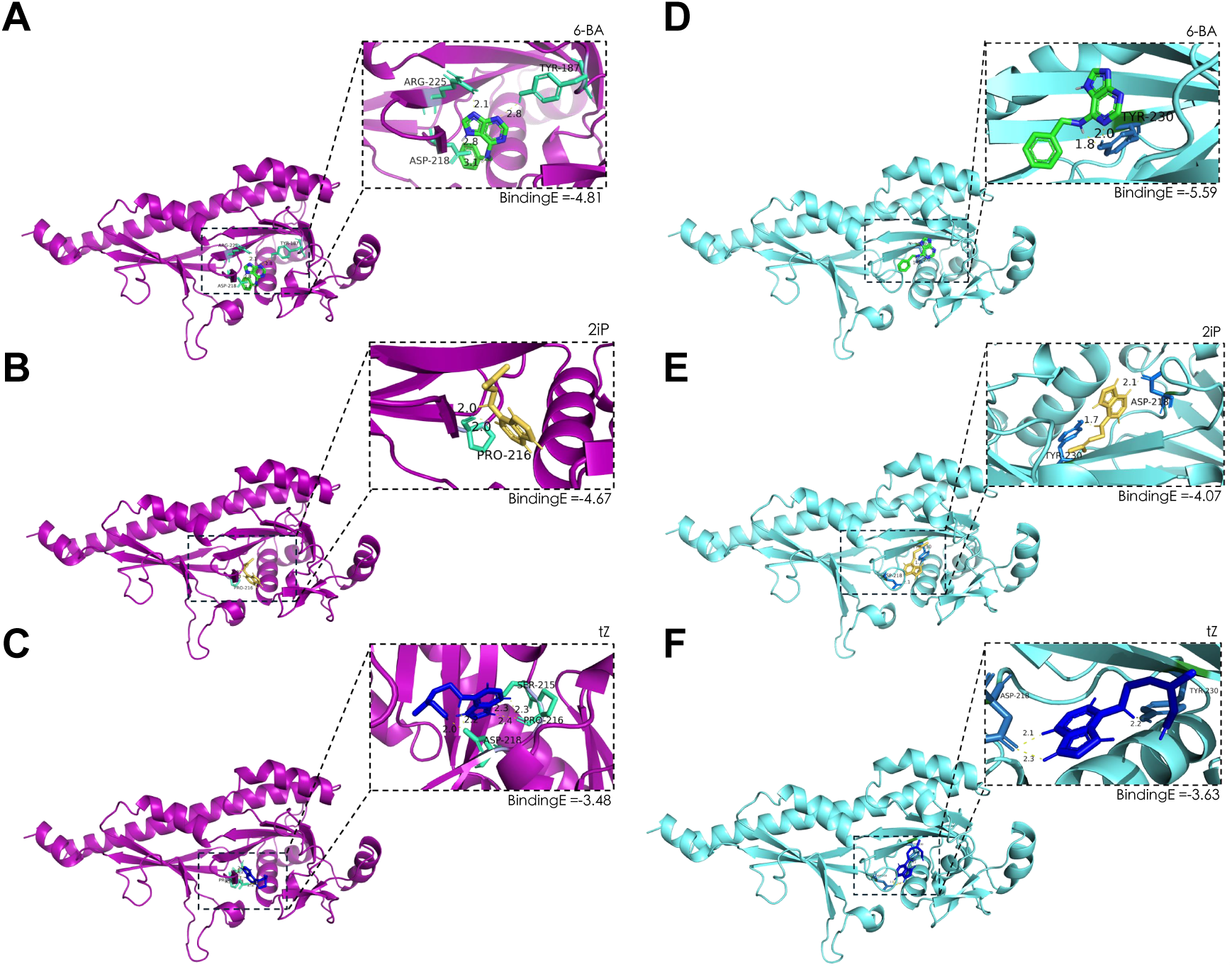
Molecular docking results of MpHK1^CD^ and MpHK2^CD^ with 6-BA, 2iP, and tZ. **(A-C)** Docking of MpHK1^CD^ (Acc. BFI26674, Mp2g03050.1 aa66-340, purple) with 6-BA, 2iP, and tZ, respectively. **(D-F)** Docking of MpHK2^CD^ (Acc. BFI16781, Mp6g00310.1 aa67-340, turquoise) with 6-BA, 2iP, and tZ, respectively. For each docking model, the overall protein structure is predicted by AlphaFold3, shown in cartoon representation. Ligand-binding pocket is magnified in insets. Direct H-bonds are indicated with dashed yellow lines, and bond length is labelled. Key residues in H-bond forming are indicated by distinct color. The predicted binding energies (kcal/mol) are shown below each inset (BindingE).

**Figure 6.**
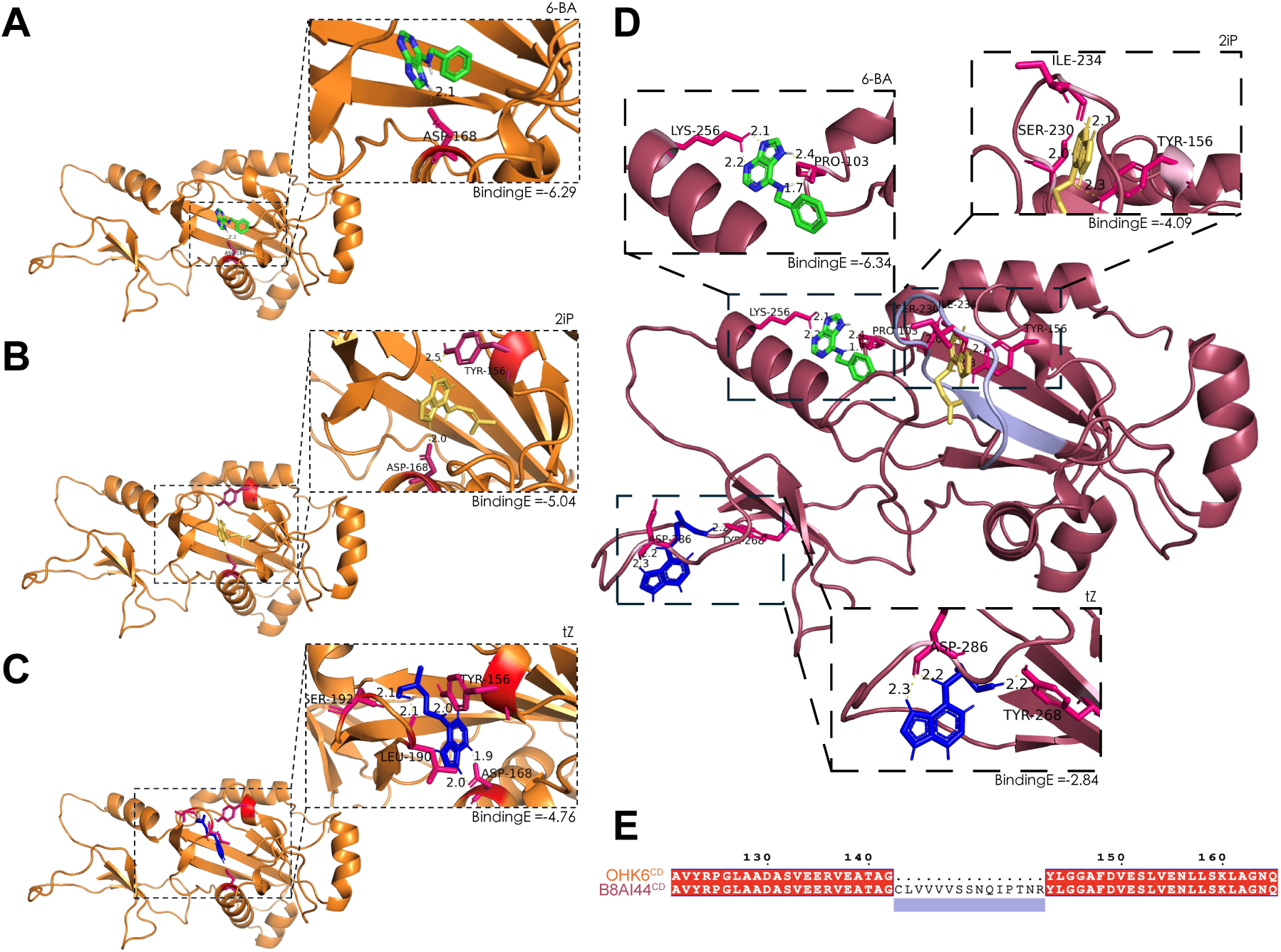
Molecular docking results of OHK6^CD^ and B8AI44^CD^ with 6-BA, 2iP, and tZ. **(A-C)** Docking of OHK6^CD^ (Acc. A1A699, Os02t0738400, aa82-294, orange) with 6-BA (**A**), 2iP (**B**), and tZ (**C**), respectively. **(D)** Docking of B8AI44^CD^ (Acc. BGIOSGA009018, aa82-309, raspberry) with 6-BA, 2iP, and tZ, respectively. The predicted binding energies (kcal/mol) are shown below each inset (BindingE). **(E)** The excess 15aa segment of B8AI44^CD^ to its homolog OHK6^CD^ is highlighted in light purple in ribbon diagram, also underlined in sequence alignment (B8AI44^CD^ aa143-157).

Across all successful docking models, each docked cytokinin formed at least one direct H-bond with pocket residues, varies from 1 to 4 H-bonds. Most H-bonds were within the range of 1.7–3.1 Å, indicating stable interactions. Generally, tZ forms more H-bonds than 6-BA and 2iP in each receptor group.

Most ligand docking positions localized within the canonical binding pocket (Figures 5 and 6A-C) except for B8AI44^CD^ (Figure 6D). Structural alignment revealed that B8AI44^CD^ contained a 15 amino acids segment longer than its homolog OHK6^CD^ (Figures 6D and 6E).

Analysis of docking models revealed that aspartic acid (Asp) and tyrosine (Tyr) residues were frequently involved in H-bonds formation across successfully docked complexes (Figures 5 and 6; Supplementary Table 1). Key H-bonds forming residues were visualized using PyMOL (The PyMOL Molecular Graphics System, Version 3.0 Schrödinger, LLC.).

### Phylogenetic Analysis of Cytokinin Receptor Homologs

The molecular docking results revealed both structural variations and conservation of key residues among cytokinin receptors. This raised the question of whether such structural divergence and conservation are associated with species’ evolution. To address this, we performed a phylogenetic analysis to place these structural observations into an evolutionary context.

To this end, a phylogenetic tree was constructed using full-length amino acid sequences of cytokinin receptors from *A. thaliana* (AHK2, AHK3, and AHK4), *M. polymorpha* (MpHK1 and MpHK2), *O. sativa* (OHK6 and B8AI44) (Figure 7). The phylogenetic tree showed a clade comprising *Arabidopsis* and *Oryza* homologs, indicating high degree of sequence conservation between monocotyledonous and dicotyledonous plants. This conservation suggests that cytokinin receptors in angiosperms may share similar ligand-recognition mechanisms. Consistent with this, our MST and docking results imply that high affinity to 6-BA might be a common feature among angiosperm cytokinin receptors.

**Figure 7.**
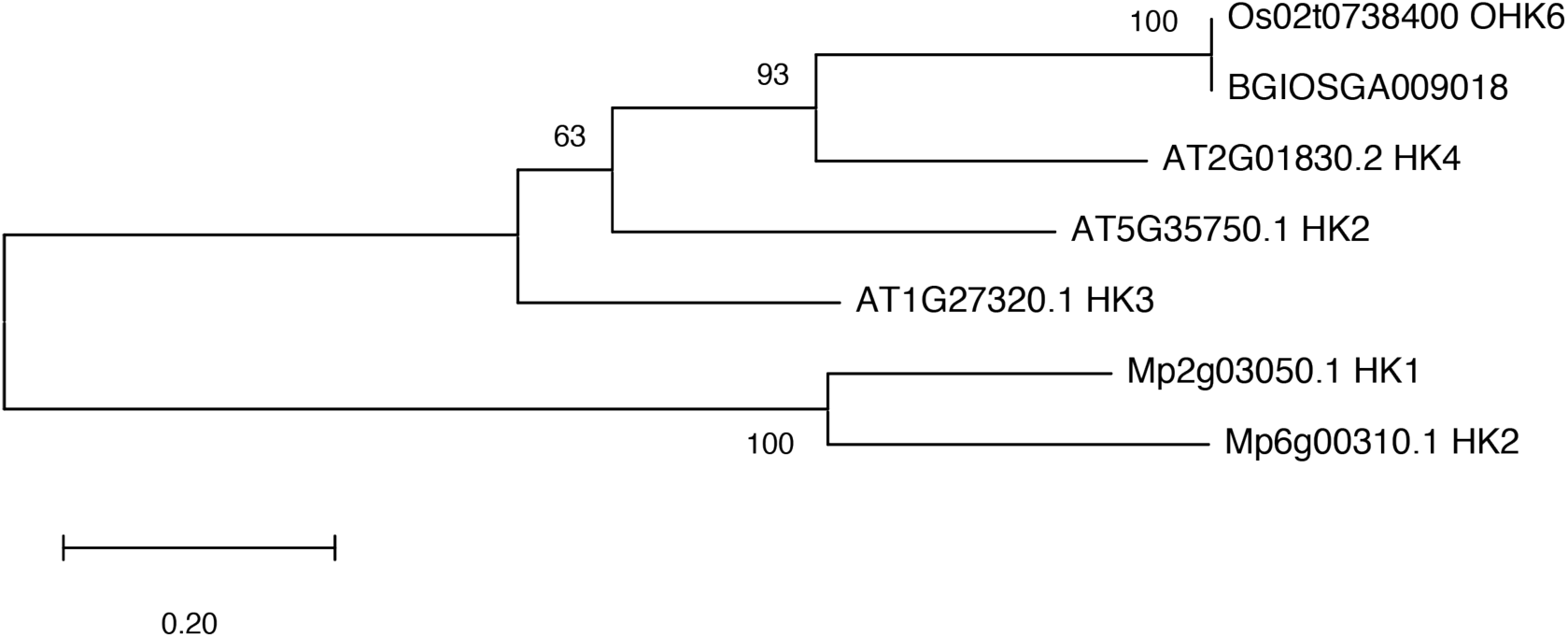
Phylogenetic relationship among cytokinin receptors from *Arabidopsis, Marchantia,* and *Oryza*. A maximum likelihood phylogenetic tree was constructed using full-length amino acid sequences of cytokinin receptors from *A. thaliana* (Acc. AHK2 – Q9C5U1, AHK3 – Q9C5U2, AHK4 – Q9C5U0),= *M. polymorpha* (Acc. MpHK1 – BFI26674, MpHK2 – BFI16781), and *O. sativa* (Acc. OHK6 – A1A699, BGIOSGA009018 – B8AI44). Bootstrap support values (percentage) are shown at the nodes, and the scale bar indicates evolutionary distance, which is number of amino acid substitutions per site.

In contrast, *Marchantia* formed a distinct clade (Figure 7), consistent with its position as a basal land plant and the early divergence in land plants (Aki et al., 2019). Despite differences in pocket structure and binding energy observed in docking, *Marchantia* homologs retain a cytokinin perception system similar to that of higher plants and can respond to exogenous cytokinins (Aki et al., 2019). This suggests that the fundamental cytokinin signaling mechanism was established early in land plant evolution and has been largely conserved.

Notably, the branch distance between OHK6^CD^ (Os02t0738400) and B8AI44^CD^ (BGIOSGA009018) was nearly zero, indicating almost identical sequences (Figure 7). However, docking analysis highlighted minor sequence differences in the binding pocket that may lead to significant variations in ligand binding behavior (Figure 6D). This observation underscores even subtle sequence divergence can affect cytokinin receptor–ligand interactions.

## DISCUSSION

### For which compound does AHK4 have a stronger binding affinity: 6-BA or 2iP?

Our MST analysis demonstrated that AHK4^CD^ exhibits a higher affinity for 6-BA than for 2iP. The 6×His-GFP control group confirms specific binding between the target AHK4^CD^ and the cytokinin ligands. However, this result contrasts with the findings by Romanov et al. (2006), who reported a higher affinity of AHK4 for 2iP (Kd_6-BA_AHK4_ = 300 nM; Kd_2iP_AHK4_ = 17 nM). A major difference between the two studies lies in the experimental systems. Romanov et al. used living transgenic *E. coli* engineered to be cytokinin-sensitive (Romanov et al., 2005; Romanov et al., 2006), whereas we employed an *in vitro* system using purified receptor proteins. This *in vitro* approach eliminates potential interference from cellular components and downstream signaling events, thereby allowing a direct assessment of the intrinsic binding affinity between the receptor and ligands. Nevertheless, accessory proteins or other *in vivo* factors, as suggested by Romanov et al. (2006), may modulate ligand-binding preferences in living systems. Thus, further validation in transgenic plants and analyses of downstream signaling pathways are necessary to establish how different cytokinins affect receptor function under physiological conditions.

The molecular basis underlying the stronger binding of 6-BA to AHK4^CD^ remains unclear. One hypothesis involves differential ligand stability. However, Hart et al. (2016) reported no significant differences in the stability of 6-BA and 2iP in aqueous environment. Structural factors are therefore more likely to contribute. Although the AHK4^CD^ crystal structures complexed with cytokinins indicate that the binding pocket can accommodate various side chains without major conformational rearrangements, subtle differences in hydrogen-bonding networks or hydrophobic interactions may favor 6-BA binding (Hothorn et al., 2011).

### Limitation of Molecular Docking Analyses

Molecular docking consistently predicted the highest binding affinity for 6-BA, as reflected by the most negative binding energies across all tested receptors. This trend aligns with the MST results, reinforcing the conclusion that 6-BA has stronger receptor interactions than other tested cytokinins.

The docking analysis also identified aspartic acid (Asp) and tyrosine (Tyr) as common residues frequently involved in ligand binding. This observation is consistent with key residues reported in the crystal structures of AHK4^CD^ (Hothorn et al., 2011; PDB: 3T4K, 3T4J), suggesting functional conservation among cytokinin receptors. However, the spatial positions of these residues varied across receptors, being located either within or at the periphery of the predicted binding pocket. Such positional variations may reflect interspecific structural plasticity, potentially shaped by evolutionary divergence.

Notably, B8AI44^CD^ consistently docked all three tested cytokinins in a peripheral binding region rather than in the common pocket. We infer that this might result from the presence of an additional 15 amino acid insertion (Figures 6D and 6E). According to structural alignment, it is positioned at the pocket entrance and may sterically hinder ligand access.

Furthermore, the number of H-bonds found in docking models was not strictly proportional to the predicted binding energy. Theoretically, a higher number of H-bonds should correspond to stronger binding. However, AutoDock’s scoring function estimates binding energy (i.e., Gibbs free energy of binding) by incorporating van der Waals forces, desolvation energy, torsional entropy, and electrostatic effects, which are not directly visualized in figures (Hill and Reilly, 2015). Moreover, the docking calculations do not incorporate water-mediated hydrogen bonds, making these predictions only an approximation of physiological binding.

In summary, randomized conformational searches by AutoDock combined with potential inaccuracies in AlphaFold3 structural predictions may produce compounded errors. Additionally, docking remains a static computational approach, unable to account for conformational flexibility or water-mediated interactions.

### Future Perspectives

In conclusion, this study demonstrates the feasibility of using MST as an *in vitro* approach to assess the binding affinity of cytokinin receptors to different ligands. Nevertheless, several limitations remain. Since cytokinin signaling *in vivo* plays a crucial role in plant responses, the *in vitro* system cannot fully recapitulate the complexity of signal transduction and metabolic regulation. To establish a more comprehensive affinity trend, additional MST assays should be performed with a broader range of cytokinins, including tZ, kinetin, and thidiazuron (synthetic phenylurea-derived cytokinin) or more (Romanov et al., 2006; Hothorn et al., 2011).

Furthermore, determining the crystal structure of receptor–ligand complexes will be essential to reveal precise binding patterns and to experimentally determine the binding energetics to validate computational predictions. Such structural information would allow the identification of all key residues involved in ligand recognition, including water-mediated H-bonds, providing a more comprehensive understanding than computational docking prediction. Based on these structural insights, site-directed mutagenesis experiments should be conducted to verify the functional significance of these key residues, thereby bridging structural predictions with receptor activity.

In summary, by providing quantitative insights into cytokinin receptor–ligand interactions and linking them with structural characteristics, this study lays a foundation for exploring the molecular basis of cytokinin selectivity. Such knowledge will facilitate the identification or design of more efficient and selective cytokinins, offering potential applications in crop improvement and advancing fundamental research on cytokinin signaling.

## ACKNOWLEDGEMENTS

This work was supported by the grants from National Natural Science Foundation of China (32388201; 31525004), Strategic Priority Research Program of the Chinese Academy of Sciences (XDB27030101), and the New Cornerstone Science Foundation through the XPLORER PRIZE to J.-W.W..

## AUTHOR CONTRIBUTIONS

X.B., C.-M.Z., and J.-W.W. conceived and drafted the manuscript. X.B. performed most of the experiments and analyses. C.-M.Z., and J.-W.W. supervised the experiments and analyses. X.B. wrote the draft and revised the manuscript together with J.-W.W..

## METHODS

### RNA Extraction and Reverse Transcription

The wild-type *A. thaliana* (Columbia-0) tissue was used for RNA extraction. Specifically, the samples were ground and mixed with TRIZOL reagent. Chloroform was then added to separate the protein, followed by centrifugation at 12,000 rpm, 4°C for 15 minutes. The supernatant was collected, and isopropanol was added for RNA precipitation. The RNA pellet was washed with 75% ethanol and dissolved in RNase-free water. DNase treatment was performed to remove any residual genomic DNA, followed by reverse transcription using a Reverse Transcriptase Kit (Thermo) to synthesize the complementary DNA (cDNA) from the extracted RNA.

### Molecular Cloning and Constructs Design

To express the fusion protein AHK4-GFP (CD, residues 126 - 395) and GFP-only in *E. coli*, the coding regions were PCR-amplified using aforementioned cDNAs as the template. The PCR products were then cloned into the double-digested pRSF-Duet vector (BamHI and SalI) via homologous recombination using the ClonExpress^®^ MultiS Cloning Kit (Vazyme). The resultant plasmids were transformed into competent *E. coli* DH5α cells, verified by Sanger sequencing, and then into *E. coli* Rosetta (BL21) cells for protein expression.

### Protein Expression and Purification

The expression vector was transformed into *E. coli* Rosetta (BL21) cells and cultured in Luria-Bertani (LB) medium with kanamycin at 37°C until the OD_600_ reached 0.8. Protein expression was then induced with 0.2 mM IPTG at 20°C and 120 rpm for 16 hours. The cells were harvested by centrifugation (4,000 rpm for 10 minutes), and the pellet was resuspended in lysis buffer (20 mM Tris-HCl, pH 7.5, 100 mM NaCl, 10% glycerol, 1 mM MgCl₂, 1 mM PMSF, and DNase). Cell lysis was performed using a high-pressure cell disruptor at 800 bar for 3 minutes. The lysate was centrifuged at 10,000 rpm for 1 hour at 4°C, and the supernatant was incubated with Ni²⁺-NTA affinity resin. After binding, the resin was washed with wash buffer (20 mM Tris-HCl, pH 7.5, 100 mM NaCl, 10% glycerol, 5 mM MgCl₂, and 25 mM imidazole). The target protein was eluted using elution buffer (20 mM Tris-HCl, pH 7.5, 100 mM NaCl, 10% glycerol, 5 mM MgCl₂, and 250 mM imidazole). The purified protein was stored in 20 mM Tris-HCl, pH 7.5, and 100 mM NaCl at 4°C for short-term analysis. Protein purity and molecular weight were confirmed by SDS-PAGE and Coomassie Blue staining.

### MST Assay

MST analysis was performed on a Monolith NT.115 (NanoTemper, Germany). The GFP-tagged AHK4^CD^ was used as the fluorescent target at a fixed concentration of 100 nM, while the cytokinin ligands (6-BA and 2iP) were serially diluted from 5 µM to 0.000153 µM. All MST buffer contained 0.05% Tween-20 to minimize adsorption in the capillaries (20 mM Tris-HCl, pH 7.5, 100 mM NaCl, 0.05% Tween-20). The measurements were performed at 25°C with 40% MST power, and three biological replicates were conducted to generate binding curves. The MST protocol was automatically generated and executed using the MO.Control v1.6.1 software (NanoTemper, Germany). Binding and dissociation constants (Kd) were calculated using the MO.Affinity Analysis v2.2.4 software (NanoTemper, Germany) by fitting the saturation binding curve at equilibrium. GFP-only protein was used as a negative control to confirm the specificity of the interaction.

### Molecular Docking Assays

Molecular docking was performed using AutoDock 4.2 with Lamarckian genetic algorithm (Morris et al., 2009; Goodsell et al., 2020) for searching optimal binding conformations. Receptor structures of MpHK1^CD^, MpHK2^CD^, and *Oryza* homologs (OHK6^CD^ and B8AI44^CD^) were obtained from AlphaFold3 predictions (Roy & Al-Hashimi, 2024), while cytokinin ligands (6-BA, 2iP, and tZ) were retrieved from PubChem (CID: 62389, 92180, 449093). Grid boxes were initially predicted by ProteinPlus (https://proteins.plus/). For docking failures, a larger grid box was applied to cover the whole receptor protein. The lowest binding energy position was selected as the predicted binding conformations. Binding energies were recorded for all conformations as reported by AutoDock. Visualization and hydrogen-bond analysis were conducted in PyMOL 3.1.6.

### Phylogenetic Tree Analysis

Full-length amino acid sequences of cytokinin receptors from *A. thaliana* (AHK2, AHK3, and AHK4), *M. polymorpha* (MpHK1 and MpHK2), and *O. sativa* (OHK6 and B8AI44) were used for phylogenetic analysis. Sequence alignment was performed using MUSCLE in MEGA12, and the phylogenetic tree was constructed using the Maximum Likelihood method (Kumar et al., 2024). Bootstrap support was calculated from 500 replicates, and the resulting percentages are indicated at the corresponding nodes. Accession numbers are listed in the figure legend (Acc.).

## REFERENCES

1. Aki, S.S. et al. (2019) ‘Cytokinin signaling is essential for organ formation in *Marchantia polymorpha*’, Plant and Cell Physiology, 60(8), pp. 1842–1854. doi:10.1093/pcp/pcz100.

2. Bauer, J. et al. (2013) ‘Structure–function analysis of *Arabidopsis thaliana* histidine kinase ahk5 bound to its cognate phosphotransfer protein AHP1’, Molecular Plant, 6(3), pp. 959–970. doi:10.1093/mp/sss126.

3. El-Showk, S., Ruonala, R. and Helariutta, Y. (2013) ‘Crossing paths: Cytokinin signalling and Crosstalk’, Development, 140(7), pp. 1373–1383. doi:10.1242/dev.086371.

4. Goodsell, D.S., et al. (2020) ‘The AutoDock suite at 30’, Protein Science, 30(1), pp. 31–43. doi:10.1002/pro.3934.

5. Hart, D.S. et al. (2016) ‘Stability of adenine-based cytokinins in aqueous solution’, In Vitro Cellular & Developmental Biology - Plant, 52(1), pp. 1–9. doi:10.1007/s11627-015-9734-5.

6. Hill, A.D. and Reilly, P.J. (2015) ‘Scoring functions for AutoDock’, In: T. Lütteke and M. Frank, eds. Glycoinformatics. Methods in Molecular Biology, vol. 1273. New York: Springer, pp.467–474.

7. Hnatuszko-Konka, K. et al. (2021) ‘Cytokinin signaling and de novo shoot organogenesis’, Genes, 12(2), p. 265. doi:10.3390/genes12020265.

8. Hothorn, M., Dabi, T. and Chory, J. (2011) ‘Structural basis for cytokinin recognition by *Arabidopsis thaliana* histidine kinase 4’, Nature Chemical Biology, 7(11), pp. 766–768. doi:10.1038/nchembio.667.

9. Jerabek-Willemsen, M. et al. (2014) ‘Microscale thermophoresis: Interaction analysis and beyond’, Journal of Molecular Structure, 1077, pp. 101–113. doi:10.1016/j.molstruc.2014.03.009.

10. Kieber, J.J. and Schaller, G.E. (2018) ‘Cytokinin signaling in plant development’, Development, 145(4). doi:10.1242/dev.149344.

11. Kieffer, M., Neve, J. and Kepinski, S. (2010) ‘Defining auxin response contexts in plant development’, Current Opinion in Plant Biology, 13(1), pp. 12–20. doi:10.1016/j.pbi.2009.10.006.

12. Kumar, S. et al. (2024) ‘MEGA12: Molecular evolutionary genetic analysis version 12 for adaptive and Green Computing’, Molecular Biology and Evolution, 41(12). doi:10.1093/molbev/msae263.

13. Morris, G.M. et al. (2009) ‘AutoDock4 and AutoDockTools4: Automated docking with selective receptor flexibility’, Journal of Computational Chemistry, 30(16), pp. 2785–2791. doi:10.1002/jcc.21256.

14. Romanov, G.A. et al. (2005) ‘A live cell hormone-binding assay on transgenic bacteria expressing a eukaryotic receptor protein’, Analytical Biochemistry, 347(1), pp. 129–134. doi:10.1016/j.ab.2005.09.012.

15. Romanov, G.A., Lomin, S.N. and Schmülling, T. (2006) ‘Biochemical characteristics and ligand-binding properties of *Arabidopsis* cytokinin receptor AHK3 compared to CRE1/AHK4 as revealed by a direct binding assay’, Journal of Experimental Botany, 57(15), pp. 4051–4058. doi:10.1093/jxb/erl179.

16. Roy, R. and Al-Hashimi, H.M. (2024) ‘AlphaFold3 takes a step toward decoding molecular behavior and biological computation’, Nature Structural & Molecular Biology, 31(7), pp. 997– 1000. doi:10.1038/s41594-024-01350-2.

17. Wulfetange, K. et al. (2011) ‘The cytokinin receptors of *Arabidopsis* are located mainly to the endoplasmic reticulum’, Plant Physiology, 156(4), pp. 1808–1818. doi:10.1104/pp.111.180539.

18. Yamada, H. et al. (2001) ‘The *Arabidopsis* AHK4 histidine kinase is a cytokinin-binding receptor that transduces cytokinin signals across the membrane’, Plant and Cell Physiology, 42(9), pp. 1017–1023. doi:10.1093/pcp/pce127.

